# Distinct laminar origins of high-gamma and low-frequency ECoG signals revealed by optogenetics

**DOI:** 10.1101/2025.10.02.680142

**Authors:** Pierre-Marie Garderes, Daniel E. Feldman, Kristofer E. Bouchard

## Abstract

Electrocorticography (ECoG) provides a high-spatiotemporal-resolution measure of cortical activity (cortical surface electrical potentials, CSEPs) in humans and animals. The CSEP high-gamma band (Hγ, 65–170 Hz) correlates with neuronal firing rates at the columnar spatial scale and is widely used as a biomarker of local activity. Whether Hγ reports all stages of columnar processing, intermediate processing in L2/3 (close to the ECoG electrode), or the main columnar output in L5, is unknown. We disentangled the laminar origins of Hγ and other ECoG bands by optogenetically suppressing L2/3 or L5 pyramidal cells during micro-ECoG recording in mouse somatosensory cortex. Whisker deflections evoked transient, topographically localized CSEPs. L5 optogenetic suppression most strongly reduced 65-450 Hz (Hγ-uHγ) bands in sensory-evoked ECoG signals, whereas L2/3 suppression most strongly reduced 4-30 Hz (θ-β) bands. Thus, different CSEP frequency bands reflect layer-specific activity and are biomarkers of distinct stages of columnar processing.

**Significance Statement:** Electrocorticography (ECoG) is widely used as a mesoscale measure of cortical activity in human and animals, providing a unique methodological from basic neuroscience discovery to understanding the human brain in health and disease. However, the cell types and laminar sources that generate ECoG signals are unknown, which impedes interpretation of ECoG findings in both the clinic and basic research. We disentangled the laminar origins of ECoG frequency bands by optogenetically suppressing L2/3 or L5 pyramidal cells during ECoG recording in mouse somatosensory cortex. We found that L5 optogenetic suppression most strongly reduced high frequencies whereas L2/3 suppression most strongly reduced lower frequencies. Thus, different ECoG frequency bands reflect layer-specific activity and are biomarkers of distinct stages of columnar processing.

## Introduction

Electrocorticography (ECoG) is widely used as a mesoscale measure of cortical activity in human clinical settings and for basic research in animals^1–10^. In micro-ECoG (µECoG), small, closely spaced surface electrodes measure cortical surface electrical potentials (CSEPs) with high spatiotemporal resolution^11^. CSEPs contain both periodic and aperiodic signal components with energy across multiple frequencies^12^. Among these, high-gamma (Hγ: 65-170 Hz) is widely used as a signature of spatially localized neuronal activity in both humans^1,4,5,7,13–16^ and animals^6,9,14,17–21^. However, the cell types and laminar sources that generate Hγ and other frequency components of CSEPs are unknown, which impedes interpretation of ECoG findings in both the clinic and basic research.

How different layers and cell types contribute to CSEP magnitude and frequency depends on biophysical factors for each cell type including distance to the surface electrode, number of cells of a given type, size and alignment of each cell’s electrical dipole, and population synchrony^2,22–24^. CSEPs originate from a linear superposition of all transmembrane currents within a volume underneath the electrode^2^. Classically, low frequencies (4-30 Hz) are primarily thought to reflect summed synaptic potentials that are the input to neuronal populations, while higher frequency ranges (particularly Hγ) are thought to reflect action potentials (spike rate) across all neurons in the volume beneath the electrode^8,9,14,18,25,26^. However, whether high-frequency CSEPs reflect the firing of all neurons in the entire underlying cortical column, or specific neuronal populations in specific layers, is vigorously debated^14,17^. In the canonical cortical microcircuit, information is processed across cortical layers in sequence from layer 4 (the thalamo-recipient input layer in sensory cortex) to L2/3 and then to L5, which is the major source of descending columnar output. If different CSEP frequency bands map to specific layers and cellular signals, these distinct CSEP bands could be used as biomarkers of columnar processing steps, providing a powerful new approach to dissect cortical computations^14,18,26^.

A recent study used simulations of a biophysically detailed model of a single cortical column from somatosensory (S1) cortex^27^ to predict the laminar contributions to different frequency bands within sensory evoked CSEPs^17^. Activating thalamocortical inputs within the model evoked a simulated ECoG signal that quantitatively resembled actual sensory-evoked CSEPs from primary sensory areas^17^. Numerical estimation within the model indicated that total transmembrane currents in L5, originating from all cell types and segments in that layer, contributed most of the high-gamma and ultra high-gamma frequency (70-450 Hz, Hγ-uHγ) signal in the CSEP, while currents in L2/3 contributed substantially less, despite being closer to the surface electrode. Because most segments in L5 belong to local pyramidal (PYR) cells, this modeling study predicts, surprisingly, that L5 pyramidal cell spiking is the largest source of Hγ in sensory-evoked CSEPs. If this is true, Hγ would reflect the final L5 output of each cortical column, while L2/3 activity (which represents an intermediate stage of columnar processing), could be more associated with a different CSEP frequency band.

Whether this preferential L5 origin of Hγ is true *in vivo* is unknown, in part because the prior simulation is single-column and thus lacks cross-columnar connections to L2/3 and other layers. Experimental studies that have sought to correlate CSEP frequency bands with neural activity in different layers have produced highly divergent findings, including that high-frequency CSEPs correlate with neural activity across all cortical layers^8^; that CSEPs mostly reflect current-source density in superficial layers^14,28^; that intracortical gamma, high-gamma, and spiking are dissociable^14,25^, and that L5 dendritic Ca2+ spikes and L1 single-unit action potentials may be observed in CSEPs^11,29^. Thus, the laminar origin of CSEP frequency bands remains fundamentally unresolved.

We took a different, causal approach using *in vivo* optogenetics to test whether L2/3 and L5 pyramidal neuron spiking contribute differentially to Hγ and other CSEP frequency bands. We recorded whisker-evoked CSEPs from mouse S1 with high-density µECoG arrays. We used the inhibitory opsin stGtACR2^30^ to optogenetically suppress spiking of either L2/3 or L5 pyramidal cells, using Drd3-Cre and Rbp4-Cre mice, which target L2/3 and L5 pyramidal cells, respectively^31–33^. We found clear evidence that Hγ and uHγ bands in whisker-evoked CSEPs predominantly originate from L5 activity, while low-frequency (θ and ß) bands preferentially originate from L2/3. Thus, different frequency components of CSEPs have distinct laminar contributions, with Hγ and uHγ being dominated by columnar output from L5. The finding that different CSEP components provide biomarkers of specific laminar activity greatly enhances the interpretability of ECoG for clinical and basic neuroscience.

## Results

### High-gamma ECoG activity in whisker S1 is somatotopically organized

To characterize sensory-evoked CSEPs in mouse S1, we recorded with high-density µECoG arrays in head-fixed, lightly anesthetized Drd3-Cre or Rbp4-Cre mice as 9 whiskers were independently deflected in a 3×3 array (**Fig. 1a**)^34^. For each µECoG channel, we used wavelet decomposition to obtain the time-frequency representation of the evoked CSEP (**Fig. 1c-e**). Stimulus-evoked power in each wavelet-defined frequency was z-scored to the pre-stimulus baseline, which enables evoked activity to be shown clearly relative to the inherent ∼1/f^a^ power distribution of the ECoG signal. We annotate six physiologically relevant frequency bands: θ (4-8 Hz), ß (10-27 Hz), γ (30-57 Hz), Hγ (65-170 Hz), uHγ (190-450 Hz) and MUA (500-1500 Hz)^2,17^. Mice expressed the inhibitory opsin stGtACR2 in either L2/3 PYR cells, L5 PYR cells, or not at all (see below).

**Figure 1.**
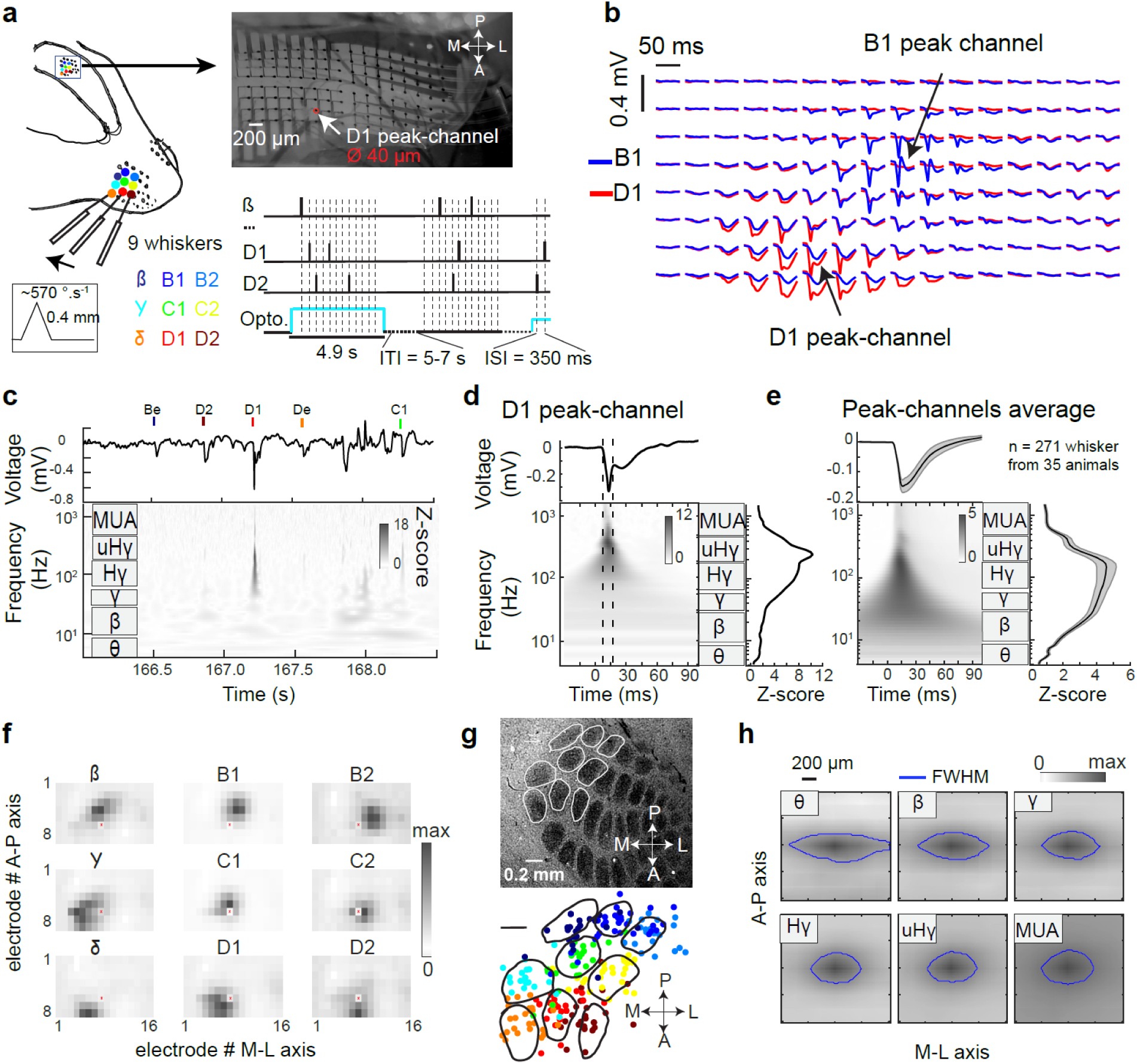
High-gamma ECoG activity in whisker S1 is somatotopically organized. **a**, Experimental approach, showing the 8 x 16 µECoG grid on the surface of S1 (pitch, 200 µm; electrode diameter, 40 µm) while whisker deflections are applied on 9 contralateral whiskers. Lower left, whisker deflection amplitude and velocity. Bottom right, two illustrative trials showing optogenetic light and whisker stimuli (at dashed lines, chosen randomly among the 9 whiskers). **b**: Mean voltage traces for all 128 electrodes for stimulation of whisker B1 (blue) or D1 (red), for one recording session. Stimulus starts at 10 ms in each trace. **c**, Raw voltage trace and time-frequency representation (z-scored to baseline epoch) in one portion of an example trial, recorded on the D1 peak-channel. **d**, Mean D1 whisker-evoked activity on the D1 peak-channel from one experiment. Right, mean evoked activity in the frequency domain, measured in the response window (dashes). **e**, Grand averaged evoked activity across all n = 271 responsive peak-channels for their columnar whisker, from 35 recording sessions. Conventions as in (**d**). Error shading is 95% confidence interval. **f**, Somatotopic structure of Hγ activity (z-score relative to baseline) on the ECoG grid for one example recording. Each subpanel is activity evoked by the labeled whisker. Each pixel is one electrode within the grid. The red dot is a defective channel. **g**, Relative spatial location of whisker-evoked Hγ peaks on the ECoG grid. Top, column boundaries from cytochrome oxidase stain for one example mouse. Bottom, location of high gamma peak for each of the 9 color coded whiskers from all sessions. Same color code as in (**a**). 8 points that were spatial outliers are omitted (see all points and details in **Supplementary Figure 1**). Barrel outlines from the example case are overlaid for illustration, to scale and manually aligned in X-Y. **h**, Mean spatial extent of whisker evoked activity on the grid (grayscale), separated by frequency band. Responses are aligned around the most responsive electrode in the indicated frequency band, and interpolated at 10× resolution. Blue line is full width at half max (FWHM).

We first analyzed the topographic organization of evoked CSEPs within whisker S1, examining trials with no optogenetic light. Single whisker stimuli evoked sensory responses at a localized subset of ECoG channels in S1 (**Fig. 1b**). For each whisker, we identified the ECoG channel with the strongest evoked response, termed that whisker’s peak-channel (**Fig. 1b**, see **Methods**). Only peak-channels with substantial responses were analyzed (n = 271/324; average Z-score >1). On average, sensory-evoked responses had power in all frequency bands, with the strongest response in Hγ, peaking 15 ± 0.2 ms (mean ± SEM) post-stimulus, consistent with the known latency of spiking responses in S1 (10-30 ms across layers)^35,36^. Subsequent analysis of effects across frequencies at a specific time examined CSEPs within a ±5 ms window around the peak of the evoked response (e.g., **Fig. 1d,e**).

Whisker-evoked Hγ CSEP signals were somatotopically organized in S1 and aligned with the known whisker map (**Fig. 1f-h**). Each whisker evoked Hγ activity in a local cluster of electrodes whose location varied with whisker identity (see example in **Fig. 1f**). When peak Hγ channel location was localized relative to whisker barrel boundaries (obtained from post-hoc cytochrome oxidase staining), the majority of peak responses (n = 128/225, localization from 25 sessions with the same nine whiskers) were located within the column boundaries for the corresponding whisker (**Fig. 1g**, see **Supplementary Figure 1** for the localization of all peak-channels). To estimate spatial spread of responses we calculated the mean CSEP activity centered on the peak-channel for each frequency band. The smallest spread was found in the Hγ (0.43 ± 0.01 mm^2^, mean area at half maximum ± SEM) and uHγ bands (0.44 ± 0.03 mm^2^). This corresponds to a circular radius of 370 µm, similar to the radius of whisker-evoked spiking in L2/3 PYR cells from 2-photon imaging (∼350 µm)^37^. Spread was significantly broader in the θ, ß and MUA bands (**Fig. 1h**, p<0.01, Friedman post-hoc test, n = 271 peak-channels) (**Supplementary Figure 2**). The spatial spread was anisotropic, extending further along the medial-lateral (M-L) axis, consistent with higher horizontal connectivity within each whisker row^38,39^. This anisotropic spread was qualitatively more pronounced at lower frequencies (θ & ß) than higher frequencies (Hγ & uHγ). Thus, like in rat auditory cortex^17^, stimulus-evoked CSEPs recorded by µECoG in wS1 are localized to approximately a single functional column and topographically organized, with activity centered on the cortical column corresponding to the deflected whisker.

### Optogenetic suppression of L2/3 versus L5 pyramidal cells

To test the contribution of L2/3 or L5 PYR neurons to whisker-evoked CSEPs, we optogenetically suppressed spiking of L2/3 or L5 PYR cells using the inhibitory opsin stGtACR2. Cre-dependent expression of stGtACR2 was achieved by injection of AAV-hSyn1-SIO-stGtACR2-FusionRed into S1 of Rpb4-Cre mice or Drd3-Cre mice, which target L5 and L2/3 PYR cells, respectively^31,32^. After 14-21 days, we performed *in vivo* neurophysiology experiments. *Post-hoc* histological analysis confirmed dense expression of stGtACR2 in L5 PYR cells in Rpb4-Cre mice, with no expression in other layers (example mouse in **Fig. 2a**, same pattern observed in all mice). In Drd3-Cre mice, L2/3 PYR cells always expressed stGtACR2, but L5 expression was also variably observed in what appeared to be thin-tufted L5 PYR cells in some mice and columns. Thus, some Drd3-Cre mice exhibited selective and dense expression in L2/3 PYR cells (**Fig. 2b, left**), while others also had expression in L5 PYR cells (**Fig. 2b, right**).

**Figure 2.**
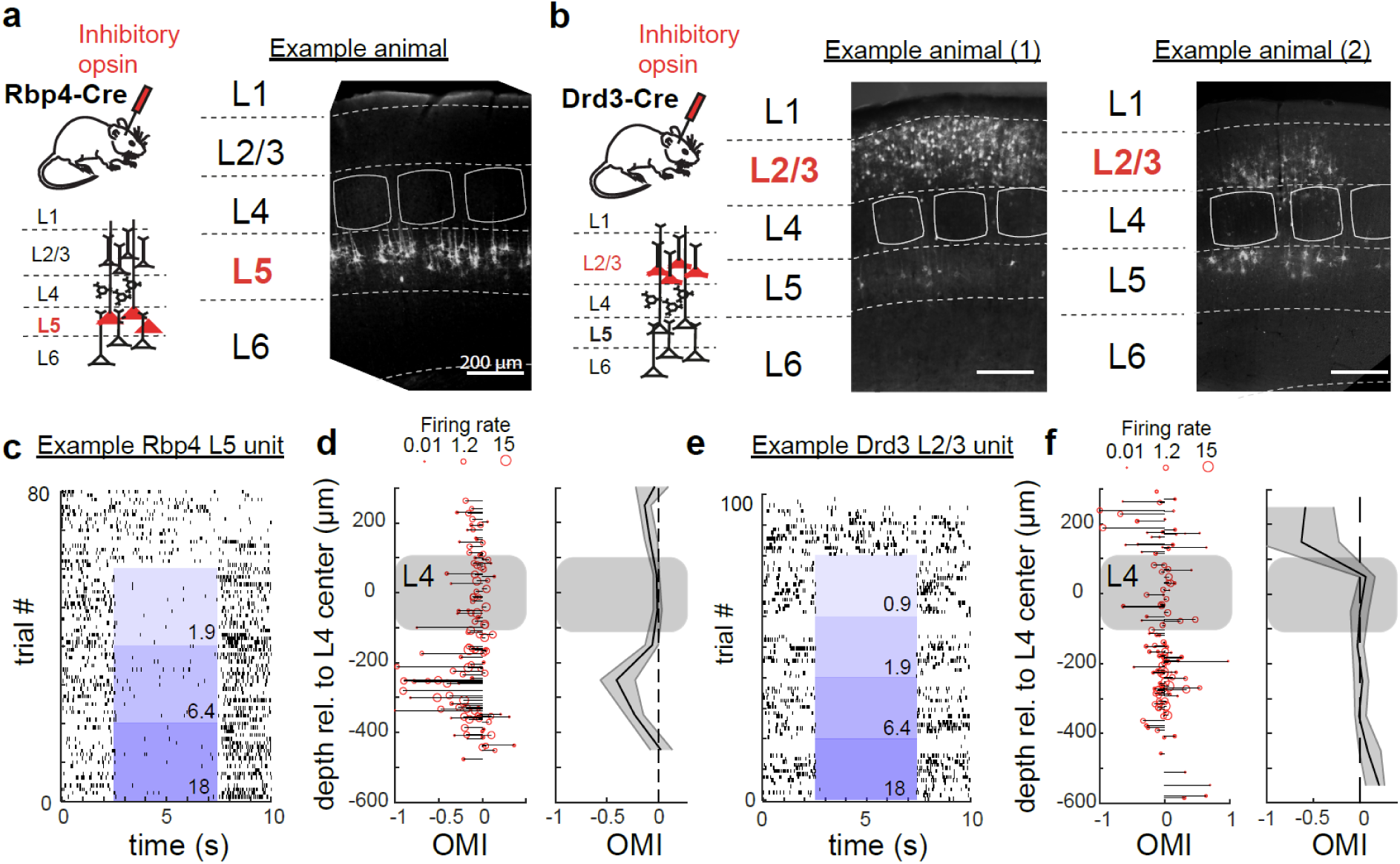
Optogenetic suppression of L2/3 versus L5 pyramidal cells. **a-b**, Histological sections from three mice, showing endogenous stGtACR2-FusionRed signal. The Rpb4-Cre mouse shows typical L5 expression (a), and the two example Drd3-Cre mice (b) were chosen to illustrate more selective (1) and less selective (2) expression in L2/3. **c,** Spike raster from a L5 multi-unit displaying optogenetic inhibition in a Rpb4-Cre mouse. Blue shows light at increasing light intensities (darker blues), with numbers indicating light power in mW. **d**, Population modulation of multi-unit activity within Rpb4-Cre mice (5 mice and penetrations, 128 units). Left, Optogenetic modulation of firing rate (OMI) for each unit across penetrations, as a function of cortical depth. Right, Average across units pooled in 100 µm bins. Shading is 95% confidence interval. **e,** Same as (c) but for a L2/3 multi-unit in a Drd3-Cre mouse. **f,** Same as (d) but for L2/3 multi-units in Drd3-Cre mice.

To test whether stGtACR2 could achieve layer-specific inhibition of L2/3 or L5 PYR cells in Drd3-Cre and Rpb4-Cre mice, we used a 32-channel laminar polytrode (spanning L2/3 to L5) to extracellularly record neuronal units *in vivo*. Spontaneous and whisker-evoked spiking were monitored, and interleaved trials contained either optogenetic light (at intensities from 0.9, 1.9, 3.5, 6.4, 11.0, or 18.0 mW) or no light (0 mW), applied to the cortical surface centered on the probe insertion site (**Fig. 2c-f**). After spike sorting^40^, we analyzed both single and multi-units and compared spike rates between light- and no-light (control) trials. The laminar location of each unit was determined by depth relative to the center of L4, identified from the pattern of whisker-evoked local field potentials on the probe (**Supplementary Figure 3**). In recordings from Rpb4-Cre mice (n = 5) with histologically confirmed L5 expression of stGtACR2 in the recorded column, we observed individual L5 units whose spiking was suppressed by light (**Fig. 2c** shows one example L5 multi-unit in response to multiple light levels). For each unit we calculated an optogenetic modulation index (OMI) that quantified firing rate changes across the full trial duration in optogenetic stimulation trials (all light levels ≥ 6.4 mW) versus interleaved no-light trials. An OMI of 0 indicates no suppression, while an OMI of -1 indicates complete suppression of spiking by light. Across all 5 Rbp4-Cre mice, L5 units (100-400 µm below L4 center) were most strongly suppressed (**Fig. 2d**, OMI = -0.219 ± 0.04, mean ± sem, n = 56 units, p<0.001 relative to OMI=0, nonparametric bootstrap test), while L2/3 units (100-400 µm above L4 center) showed a much weaker suppression (OMI = -0.079 ± 0.03, n = 25 units, p=0.016). In contrast, in Drd3-Cre mice (n = 5) that were confirmed histologically to have L2/3 expression but sparse or no L5 expression in the recorded column, L2/3 units were often suppressed by optogenetic light (an example L2/3 multi-unit is shown in **Fig. 2e**). Across these Drd3-Cre mice, average spiking was significantly suppressed for L2/3 units (**Fig. 2f**, OMI = -0.615 ± 0.17, mean across n = 18 units, p=0.006), but not for L5 units (OMI = 0.014 ± 0.02 OMI, n = 66 units, p=0.7). The minor suppression of spiking for units in non-genetically targeted lamina may reflect indirect circuit-level effects of target layer suppression. Thus, stGtACR2 suppressed mean firing rate by 22-60% in a layer-specific manner: Rpb4-Cre mice showed fairly selective optogenetic suppression of L5 spiking, while Drd3-Cre mice showed selective suppression of L2/3 spiking. This matches the histological expression patterns of stGtACR2.

### Different CSEP frequency bands are modulated by suppression of L2/3 versus L5

To test how layer-specific suppression affects CSEPs, we performed µECoG recordings while applying optogenetic light at varying levels, or no light, through the transparent µECoG array on interleaved trials. Optogenetic illumination covered multiple cortical columns with a diameter of ∼0.63 mm (full width at half maximum). These recordings were separate experiments from the intracortical recordings and came from a larger set of mice (10 Rpb4-Cre mice, 12 Drd3-Cre mice, and 3 uninjected C57BL/6 mice). Experimental variability in the light intensity and viral-mediated opsin expression at each spatial location in S1 will drive variability in optogenetic effects at different peak-channels. We therefore quantified opsin expression in L2/3 and L5 at each peak-channel location in each mouse (**Fig. 3a-b)**. To do this, we prepared tangential histological sections (parallel to the pial surface) and measured the density of FusionRed-positive neuronal somata (which marks stGtACR2 expression) in L2/3 and L5. Expression density was quantified in a hexagonal area centered on the cortical column for each peak-channel (e.g., on the anatomical D2 column for the D2 whisker’s peak-channel, **Fig. 3a**). We then classified peak-channels into 4 groups based on layer-specific opsin expression and light delivery (see **Methods** and **Supplementary Table 1**):

1. Peak-channels from mice with no stGtACR2 expression (n = 63 channels, from 3 uninjected C57Bl/6 mice, and from 3 Drd3-Cre and 2 Rbp4-Cre mice with failed stGtACR2 expression).
2. Peak-channels with selective expression in L5, but not L2/3 (n = 35 channels from 6 Rbp4-Cre mice).
3. Peak-channels with selective expression in L2/3, but not L5 (n = 25 channels from 6 Drd3-Cre mice).
4. Peak-channels with expression in both L2/3 and L5 (n = 15 channels from 4 Drd3-Cre mice)

**Figure 3.**
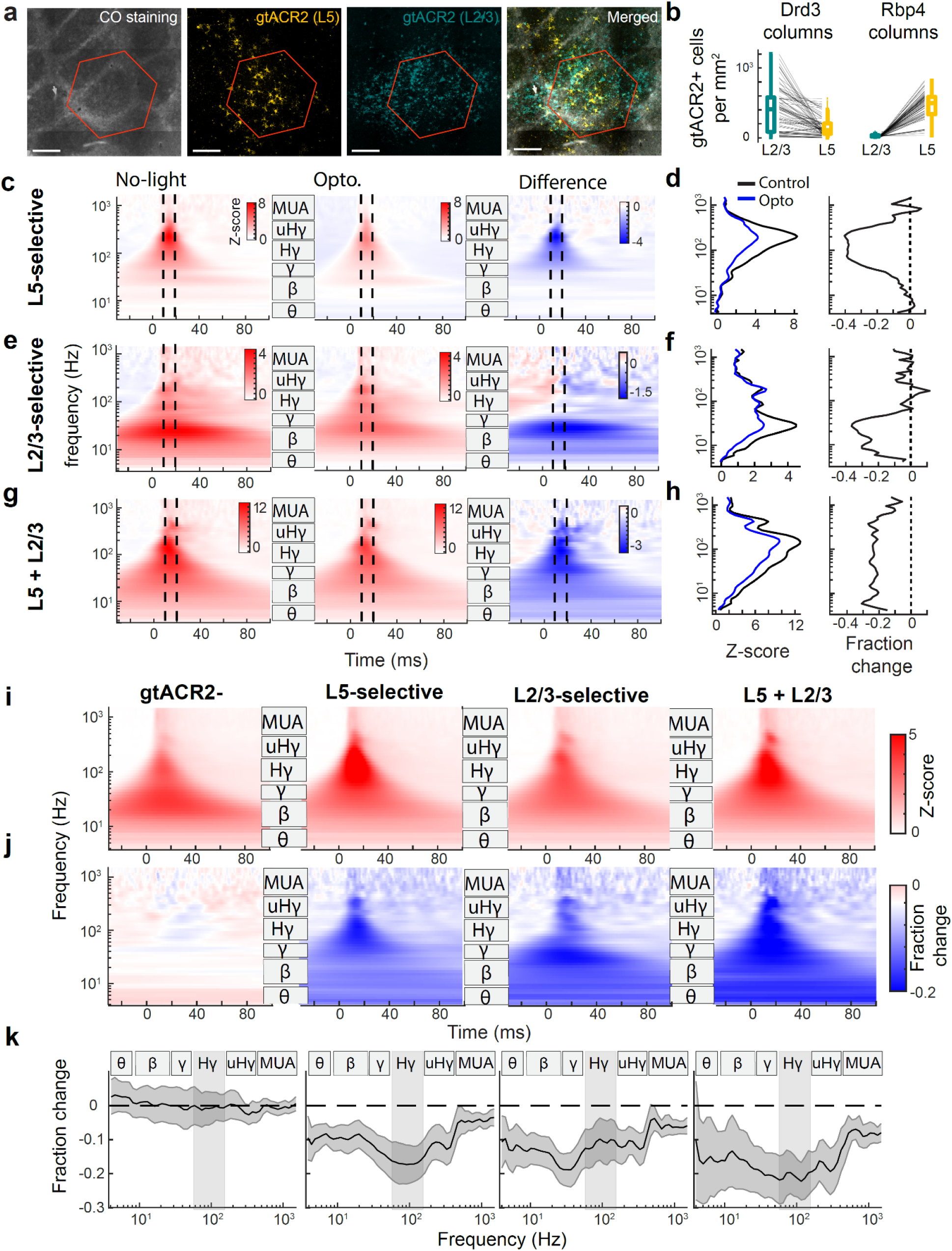
Different CSEP frequency bands are modulated by suppression of L2/3 versus L5. **a**,Tangential sections from an example Drd3-cre mouse, with hexagon for quantifying stGtACR2 expression around the D1 peak channel. FusionRed reporting stGtACR2 in L5 (gold) and L2/3 (teal) is shown relative to CO staining in L4 (black). **b**, Quantification of stGtACR2 in L23 and L5 in all cortical columns from all 17 mice with histological data. **c, e, g**, Average CSEP spectrogram for three peak-channels with selective expression in L5 (c), selective expression in L2/3 (e), and expression in both L5 and L2/3 (g). Left, spectrogram for whisker-evoked CSEP in the no-light condition (“Control”). Middle, light condition (“Opto”). Right, spectrogram difference Opto - Control. **d, f, h**, Quantification of CSEP magnitude by frequency for Control and Opto conditions, and the fraction change. These were quantified in the ±5 ms time window around peak response. **i-k,** Population average analysis. **i,** Z-scored evoked responses for all peak-channels within each group defined by stGtACR2 expression. **j**, Optogenetic suppression computed as the fraction change of the spectral power, computed from the raw wavelet magnitude as light/no-light - 1. **k**, Quantification of the fraction change in j. Shading is 95% CI across peak-channels.

We evaluated the impact of optogenetic light in these four groups by analyzing CSEPs evoked by the stimulation of the peak-channel’s whisker. Focusing first on the groups with stGtACR2 expression, **Figure 3c–h** shows example peak channels from the groups with either L5-selective, L2/3-selective, or L2/3+L5 expression. We compared the CSEPs spectrogram (z-scored to baseline activity in each frequency) in light-on trials (≥6.4 mW) versus no-light trials by plotting the difference (ΔZ-score) of these spectrograms, and the fractional change relative to no-light trials (using non-Z-scored frequency band power). The ΔZ-score and fractional change were then quantified for all frequencies. In these representative examples (**Fig. 3c–h)**, selective L5 suppression led to a reduction in Hγ and uHγ power, selective L2/3 suppression reduced low-frequency activity (θ, β, and γ bands), and combined L2/3+L5 suppression resulted in broadband attenuation across frequency bands.

We assessed the population average effect of optogenetic suppression on CSEPs within each layer-specific group (**Fig. 3i-k).** The average CSEP spectrogram across all peak-channels for the no-light condition (z-scored to baseline activity) revealed robust responses to whisker deflection across frequency bands in each group (**Fig. 3i**). Next, for each peak-channel, we computed the fractional change for each frequency relative to no-light trials and the resulting spectrograms were pooled within each group (**Fig. 3j**). We then quantified this fractional change for individual frequencies within a ±5 ms window around the peak of the evoked response (**Fig. 3k**).

We found that channels with no stGtACR2 expression had no optogenetic modulation in CSEP power across any frequency band, assessed by the fractional difference in power between light and no-light trials (0.01 ± 1.7 % opto-modulation from 4 to 1500 Hz mean ± sem across peak-channels). Channels with selective L5 expression showed the strongest suppression in the Hγ band (−16.6 ± 2.5 % from 65 to 170 Hz), but significant suppression in all frequency bands (see confidence intervals in **Fig. 3k**). Channels with selective L2/3 expression showed the most prominent suppression in the ß and γ frequency bands (−14.4 ± 2.2 % from 10 to 57 Hz), again with significant suppression in all frequency bands. Channels with combined L2/3 and L5 expression showed a broad-spectrum suppression spanning both low and high CSEP frequencies (−17.8 ± 3.6 % from 4 to 450 Hz). Thus, this population average analysis reveals that suppression of L2/3 PYR neurons most prominently suppresses low CSEP frequencies, while suppression of L5 PYR neurons preferentially suppresses Hγ CSEP activity.

### L5 vs. L2/3 preferentially contribute to high vs. low-frequency CSEP signals

The population-level results above demonstrate that optogenetic suppression of L2/3 and L5 preferentially suppress low and high-frequency CSEPs, respectively. However, we know that there is substantial variability in, e.g., the level of opsin expression, etc., which will modulate the relative magnitude of the effects. Therefore, we used a linear mixed-effects model to quantify the relative contribution of L2/3 and L5 to optogenetic suppression across CSEP frequency bands (**Fig. 4**, see Methods). Unlike the population-average approach shown in **Figure 3**, the model accounted for the quantitative level of opsin expression (determined histologically) at each peak-channel location, spatial and trial-by-trial variability in light exposure, and statistical dependence between peak-channels in the same recording session. In addition, the model included the entire dataset with histologically quantified levels of stGtACR2 expression, not just the channels that fit within the four groups of **Figure 3**. Optogenetic suppression (ΔZ-score) was modeled using two fixed effects: ß_L2/3_X_L2/3_ and ß_L5_X_L5_ where X_L2/3_ and X_L5_ represent the estimated reduction in spiking in either L2/3 or L5 PYR cells predicted from quantitative levels of opsin expression and light exposure at each peak-channel. A nested random effect (whisker identity) controlled for repeated measurements within recordings. The model predictive performance was quantified with leave-one out cross-validation.

**Figure 4:**
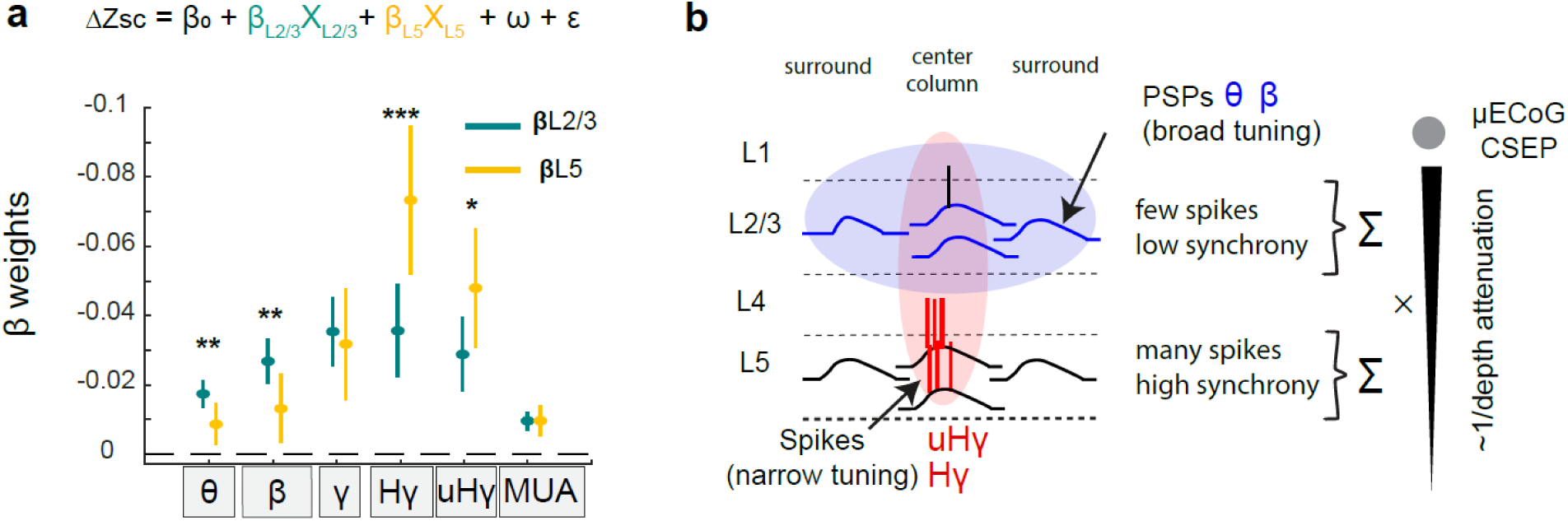
L5 vs. L2/3 preferentially contribute to high vs. low-frequency CSEP signals. **a**, Mixed effect model of CSEP suppression. We modeled suppression in each canonical frequency band based on putative spiking suppression of L5 and L2/3 PYR cells, estimated from local stGtACR2 expression and light intensity in individual peak-channels. ß weights represent contribution of each layer’s activity to the frequency band in the CSEP. Error bars represent the 95% confidence intervals. Statistical tests are conducted using a randomization procedure. **b**, Conceptual model of laminar contributions to ECoG (see discussion). Left, schematic voltage-time traces for individual sources in L2/3 and L5 center and surround columns. L5 PYR exhibits high firing rate columnar tuning and strong dipole generating Hγ at the surface. Local L5 PSPs are attenuated by depth. L2/3 PYR exhibit sparser spikes with the contribution to CSEP of the layer being dominated by the broadly tuned PSPs, integrating over longer time windows.

The model was fit independently for each of the six frequency bands (θ, β, γ, Hγ, uHγ, and MUA). When evaluated on held-out data, the models achieved significant predictive performance, with p-values <10^-16^ and coefficients of determination (R²) ranging from 0.26 to 0.42 across bands (**Supplementary Table 2)**. In this model formulation, negative ß values represent a suppression of the frequency band in response to light. In all frequency bands, the fitted coefficients ß_L23_ and ß_L5_ were significantly below 0, indicating that suppressing L2/3 or L5 activity both significantly reduced CSEP power in that frequency band (**Fig. 4**). However, we found a significantly higher contribution of L5 as compared to L2/3 in the Hγ band (ß_L5_ /ß_L2/3_ = 2.06; p = 0.0007, corrected for multiple comparisons, see Methods and **Supplementary Table 2**). This was also true for the uHγ band (ß_L5_ /ß_L2/3_ =1.67; p = 0.018). Interestingly, we found a significantly higher contribution of L2/3 as compared to L5 in the θ band (ß_L2/3_/ß_L5_ =2.02; p = 0.002) and β band (ß_L2/3_/ß_L5_ = 2.04; p = 0.005). Together, these results establish that L5 is the primary source of Hγ and uHγ ECoG activity (65–400 Hz), whereas L2/3 drives lower-frequency components (4–30 Hz). This demonstrates a clear laminar specialization in sensory evoked ECoG activity.

## Discussion

### Resolving the inverse problem of CSEPs with causal manipulation of targeted source neurons

Identifying the precise neuronal sources that combine to produce the CSEP is known as the ’inverse problem’ for CSEPs. Solving the inverse problem is inherently difficult because diverse combinations of neuronal sources can, in principle, produce similar surface potentials– that is, it is an ill-posed inverse problem^2,41^. Further complications arise from the proximity in time and space of activation of multiple cell types across layers, and the contribution of action potentials, synaptic potentials, dendritic calcium spikes, and other signals within and across cells. Previous efforts to identify the contributions of specific neural layers and sources to CSEPs have primarily relied on correlations between ECoG, intracortical LFP, and spiking *in vivo*^8,14,18^ or using computational simulations^17,42^. While informative, these approaches have inherent limitations. Correlation studies are limited by the near-simultaneous activation of all cortical layers, and fail to systematically account for key biophysical factors like neuronal synchrony, size of produced dipoles (due to neuronal geometry), and total number of neurons^17,22^. Computational models, while offering valuable theoretical insights and predictions, depend on simplifying assumptions that may be inaccurate, such as omitting certain cell types or even whole layers, using biophysically oversimplified neurons, or lacking cross-columnar interactions^17,42–44^. As a result, the origin of Hγ signals (which are widely studied by ECoG) and other frequency bands within CSEPs have remained unresolved.

In particular, it has been unclear whether Hγ and uHγ arise from very local sources (e.g., a single cortical column) and what is the laminar origin of these signals. To resolve this, we used causal layer-specific manipulation to determine the relative contribution of L2/3 and L5 PYR cells to sensory evoked CSEPs recorded by µECoG in mouse S1. Our results show that sensory-evoked Hγ CSEP signals are temporally brief and display sharp, column-scale topographical localization **(Fig. 1 and SFig. 1**). Using layer-specific optogenetic suppression of L2/3 or L5 PYR cells (**Fig. 2**), we found that L5 suppression preferentially decreased the high frequencies (Hγ and uHγ bands) within CSEPs, while L2/3 suppression preferentially decreased lower frequencies (θ and β). Using a mixed effects model, we found that L5 exerts twice the weight, relative to L2/3, on Hγ-uHγ CSEP bands, while L2/3 contributes twice as strongly to the θ-β bands (**Fig. 4a**). Thus, L2/3 and L5 PYR cells differentially generate low- and high-frequency CSEP components (**Figs. 3-4**). These causal experimental results validate the findings of a prior computational study that L5 PYR neurons in the local cortical column are the primary contributors to high-frequency CSEP activity^17^, and do not support alternative hypotheses that all layers contribute equally, or that L2/3 is the predominant source of high-frequency CSEPs^8,14,42–44^.

These results can be explained by the well-established neocortical principle that while both L2/3 and L5 PYR cells exhibit postsynaptic potentials (PSPs) to columnar input, L2/3 PYR neurons exhibit fewer spikes and a sparser code than L5 PYR neurons^45^, with L5 PYR cells producing more total spikes and a higher mean firing rate per neuron, including spike bursts^46^. These differences in spiking statistics are conserved across cortical areas and species^14,45,47–49^, and have important consequences for extracellular field generation. Specifically, dense spiking from L5 generates large local fields dominated by high-frequency power that propagate to the cortical surface, while sparse spiking from L2/3 generates local fields dominated by low-frequency power because of PSPs whose slow kinetics allow them to be integrated over extended time windows^2,24^. Our findings map directly onto this canonical principle (**Fig. 4b**). Suppressing L2/3 pyramidal neuron activity selectively attenuated low-frequency components (4–30 Hz) of the sensory-evoked ECoG signal, which were spatially elongated along the whisker row (**Fig. 1h**), consistent with PSPs driven by anisotropic connectivity of L2/3 cross-columnar projections^38,39^). In contrast, suppressing L5 pyramidal neuron activity selectively attenuated high-frequency components (65-450 Hz), which were sharply localized in space, consistent with synchronous firing, strong dipoles, and narrow sensory tuning of spikes in L5 neurons, and the attenuation of deep-layer PSPs at the cortical surface. Together, this conceptual model explains how L2/3 contributes predominantly to low-frequency surface signals via local, broadly tuned PSPs^50–52^, while L5 contributes to high-frequency signals via dense and synchronous spiking^46,53^. Therefore, this model offers a mechanistic bridge between canonical laminar functional properties and the frequency structure of ECoG signals.

### High-gamma activity preferentially reports cortical column output

Together, our findings indicate that Hγ activity does not reflect the simple average activity across layers, but instead more strongly reflects L5 PYR spiking activity compared to other sources. In sensory cortex, the canonical feedforward cortical microcircuit starts in L4, the main thalamic recipient layer, and progresses to L2/3 and then L5^54,55^, though actual processing is more complex^56^. As information propagates through this laminar hierarchy, L2/3 and L5 perform nonlinear integration of feedforward input with contextual information from nearby columns and additional context and expectations from top-down inputs^57–59^. The results of these columnar computations are then broadcast to the rest of the brain, shaping behavior and sensory decisions^60–62^. L2/3 cells represent an intermediate stage of intracolumnar processing and provide output to higher cortical areas, while L5 cells are the final output to cortical and subcortical targets, including higher-order thalamus, striatum, and motor-related structures in the midbrain and brainstem^63^ and are most related to perceptual decisions and motor control^60^.

As L5 spiking activity dominates Hγ in ECoG signals, Hγ provides a biomarker for the final L5 output of columnar computation. In contrast, lower frequency bands may reflect more intermediate stages of computation, e.g., in L2/3, and from input PSPs prior to cellular processing into spike trains. In S1, sensory-evoked Hγ is transient and non-oscillatory, and thus likely reflects brief periods of synchronous high firing in L5 PYRs of individual S1 columns. The transient nature of this signal is consistent with a model in which processed columnar output is not transmitted continuously but in brief bursts, which may increase effectiveness of information transmission to downstream targets^64–66^.

Hγ activity in CSEPs is associated with a variety of perceptual, behavioral and cognitive processes in animals^21,67,68^ and humans^7,10,13,69^. The finding that Hγ is a biomarker of columnar output, rather than internal columnar processing, provides an explanation for the remarkable tuning of Hγ to sensory and motor variables. In sensory systems, this includes sharp sensory tuning of Hγ in animal models^17,26^, which tracks the sharper sensory tuning of L5 neurons than L2/3 neurons^35,70,71^, and sharp tuning for speech acoustics in humans^72^. In motor areas, such as those controlling vocal tract articulators in humans^13^, sharp Hγ tuning may reflect the motor output of L5 neurons rather than preparatory activity, which may explain the success of decoding motor commands from these ECoG signals. The paradoxical observation that surface Hγ has sharp, topographically organized tuning may thus reflect its readout of finely tuned activity and precise topographical projections of L5 neurons.

### Open questions and future directions

Together, our findings indicate that Hγ ECoG activity is an effective mesoscopic readout of cortical column spiking output in L5, relative to input or internal processing. Likewise, the relative contributions of within column vs. across column PSPs in L2/3 that drive low-frequency ECoG signals remain unclear, and more detailed modeling and higher-resolution manipulations will be essential to disentangle these sources. Looking ahead, how broadly do these laminar-spectral relationships generalize across cortical regions and behavioral states? Although laminar organization is conserved throughout the neocortex, regional differences in layer thickness and cellular composition may shape how different lamina are reflected in frequency bands of CSEPs, and hence how those frequency bands map onto cortical processing.

## Conclusion

Our findings offer a new functional interpretation of distinct ECoG frequency bands as biomarkers of neural activity in different cortical layers. ECoG Hγ reflects L5 columnar output, while low frequency signals are more linked to L2/3, and hence may reflect intracolumnar processing. Thus, different frequency bands of CSEPs may be biomarkers of distinct computational stages of columnar processing. Enhancing our understanding of the sources of ECoG signals^73^, especially from high-resolution, large-scale recording devices^74,75^, could enhance performance of neuroprosthetic systems^76^ and diagnostic tools. Large scale µECoG systems that densely tile multiple cortical areas may be a valuable research tool by enabling millisecond resolution columnar-scale readout of different stages of laminar processing across cortical regions.

## Author Contributions

D.E.F. and K.E.B. conceived the study and supervised the project. P.M.G., D.E.F. and K.E.B. designed the experiments and analysis. P.M.G. conducted the experiments and performed the analysis. P.M.G. wrote the initial draft. P.M.G., D.E.F. and K.E.B. reviewed and edited the manuscript.

## Acknowledgments

This work was supported by NIH R01 NS118648 (to D.E.F. and K.E.B.), LBNL LDRD ‘Neural Systems and Machine Learning lab’ (to K.E.B.), and NIH R01 NS092367 (D.E.F.)

## Methods

### Animals

Data from 3 WT, 18 Rbp4, 17 Drd3 animals were used in this study. All animal procedures were performed in accordance with local regulations as described in the IACUC Protocol # AUP-2016-02-8351-2. (Institutional Animal Care and Use Committee). WT and Drd3 [(Tg(Drd3-cre) KI196Gsat / Gensat MMRRC (strain number 034610) JAX #)] (Gensat MMRRC, strain number 034610) mice were C57Bl6 background while Rbp4 [(Tg(Rbp4-cre) KL100Gsat / (GENSAT line # KL100))] were on Agouti/C57Bl6 mixed background. Animals used were aged 2-11 months and were either male or female.

### Surgeries

An initial viral injection was performed in Rbp4 and Drd3 mice. Mice were anesthetized with isoflurane (induction at 5%, maintenance at 2%) and placed in a stereotaxic frame over a feedback-controlled heating pad set to 37 °C. Eyes were protected from desiccation using a petroleum-based eye ointment. General analgesia was provided with Meloxicam (5 mg/kg, subcutaneous), buprenorphine (0.1 mg/kg, subcutaneous), and dexamethasone (4 mg/kg, subcutaneous). The incision area was shaved and disinfected with betadine, and lidocaine was injected along the midline under the scalp (1%, 0.1 mL, subcutaneous). A 10 mm incision was made to expose the skull. A burr hole was drilled at the coordinates of C1 or C2 whisker barrel in the left hemisphere (AP 1.4, ML -3.3 relative to bregma) using a dental drill. A beveled glass pipette with a diameter below 30 µm was inserted at a depth of 400–500 µm. Between 400–600 nL of the viral construct pAAV_hSyn1-SIO-stGtACR2-FusionRed (a gift from Ofer Yizhar; Addgene viral prep # 105677; RRID:Addgene_105677) was injected at a rate of approximately 60 nL per minute. The pipette was removed 3 minutes after the injection concluded. The skin was sutured, and the animal was placed in a recovery cage, half over a heating pad, until fully awake. Animals were monitored for three days post-surgery, with meloxicam (5 mg/kg, subcutaneous) administered once daily. Optimal transfection density and stability were achieved by diluting the viral construct in Ringer’s solution to a final concentration of 1 x 10^12^ vg/mL, and waiting 14–21 days post-infection before recording. Control animals received either sham viral injections (Drd3 and Rbp4 mice) or no injection at all (WT).

Following the incubation period, The mouse was prepared using the same protocol as for viral injection until incision. Then the skull was cleaned using a scalpel, the skin was secured away from the skull using surgical glue, and the bone was maximally thinned on the edges of a rectangular craniotomy using the dental drill. The rectangular craniotomy area measured approximately 5mm (Medio-lateral axis) by 3 mm (Antero-posterior axis) and was centered to a point 1 mm lateral to the C1 whisker barrel. A burr hole was drilled in the frontal region of the ipsilateral hemisphere to insert a 30 gauge silver-wire connected to a golden pin which would serve as reference for the electrophysiological recordings. Dental cement was used to seal a custom metal headpost in the back of the skull. The area was kept moistened with saline for at least 5 minutes to limit bleeding at bone removal. The bone flap was removed with a tweezer and the animal placed in the setup for electrophysiological recording.

### Electrophysiological recordings

The protocol for simultaneous electrophysiological recording and optogenetic inhibition has been described in detail previously^34^. Briefly, during the recording session, animals were maintained in a lightly anesthetized state (Isoflurane 0.5–1% in 0.4 L/min O₂), and placed on a feedback-controlled heating pad set to 37 °C with head-fixation. Chlorprothixene was administered (1 mg/kg, intraperitoneal) to help maintain anesthesia at a lower isoflurane level.

Neural data from ECoG were acquired using a custom-designed 128-channel µECoG grid (E128-200-8-40-HZ64x2, Neuronexus, Ann Arbor, MI, USA) connected to a 128-channel headstage (SpikeGadgets, San Francisco, CA, USA). The grid was placed on the moistened brain and carefully moved in a medio-lateral direction. Once in position, the remaining saline solution was aspirated, and the grid was gently pressed against the brain in a latero-medial movement, allowing it to bend slightly to better conform to the shape of the mouse brain, ensuring optimal contact between electrodes and the cortical surface. No additional saline was added during the rest of the ECoG recording. If needed, the grid was repositioned following a short somatotopic mapping session (20–30 deflections per whisker) to confirm proper positioning. Electrode contacts on the grid had an exposed diameter of 40 µm with a 200 µm inter-electrode pitch. Data was recorded at a sampling rate of 30 kHz using Trodes software version 2.3.3 (SpikeGadgets, San Francisco, CA, USA), downsampled to 3 kHz, and saved in NWB format.

Neural data from the laminar probe recording were collected either in separate animals or after the ECoG recording session. We used 32-channel probes (A1x32-Poly2, Neuronexus, Ann Arbor, MI, USA) and a 32-channel headstage (SpikeGadgets, San Francisco, CA, USA). Duratomy was performed locally with a 30G hypodermic needle, and the probe was inserted at a speed of <2 µm/sec. The electrodes were arranged on the probe in a staggered layout every 25 µm over a total length of 800 µm, enabling simultaneous sampling of supragranular, granular, and infragranular layers. Recording began 10 minutes after the probe was fully inserted. Shielding and grounding of the setup minimized significant 60 Hz noise in the electrophysiological recordings.

Once the recording devices were in place, optogenetic light was directed at the cortical surface, and its position adjusted to match the expression area. Light from a 473 nm LED was collected in a 1 mm diameter optical fiber using a tandem of aspheric lenses. Light from the fiber was delivered through a lens and focused on the surface to create a ∼0.63 mm diameter disk (FWHM). In optogenetic trials, light was applied for 4.9 seconds, starting 100 ms before the first whisker stimulus and ending 450 ms after the last whisker deflection in the trial. Optogenetic light intensities were randomly varied from trial to trial at one of the following values: 0 (control), 0.9, 1.9, 3.5, 6.4, 11.0, or 18 mW. In 25% of trials, light was applied without whisker stimulation.

An array of 3x3 whiskers was inserted into glass capillaries attached to individually controlled piezo actuators. Whisker deflections were designed as 12 ms back-and-forth pulse deflections. A 10th line was used as a blank stimulus, randomly intermingled with whisker pulses. Within each trial, 12 whisker pulses were randomly intermingled and delivered with a 350 ms inter-pulse interval over the total 4.9 s duration of the trials. Trials were separated by a 5 to 7 s inter-trial interval.Tactile stimuli were created through custom Igor code and delivered to the piezo actuator at a sampling frequency of 200 kHz. Onset times of whisker stimulation were recorded as TTL pulses in the electrophysiological system with 1 ms precision, allowing for later alignment of stimulus-evoked CSEPs. The inter-trial interval was varied between 5 and 8 seconds. On average, 289.3 ± 11.4 trials (mean ± SEM) were recorded per animal .

### Analysis Code

All analyses were performed using custom code in MATLAB.

### Histology

In a subset of 17 animals, 3-5 distinct points in the brain were marked with DiO to facilitate later alignment of histological slices. Animals were euthanized following the recording session. The brain was collected and immersed in 4% PFA. The next day, neocortical tissue around the craniotomy area (∼6x4 mm) was extracted, flattened between two microscope slides, and placed in a 30% sucrose solution for at least 48 hours. The brain was then frozen and sliced tangentially to the surface in 50 µm sections. Slices were stained with cytochrome oxidase (Cytochrome C from equine heart; Medix Biochemica, Finland) to reveal the position and contours of the whisker barrels in layer 4 slices.

Images of each slice were taken using a confocal microscope and re-aligned along the XY dimension with the EMTrak2 plugin^77^ via DiO labeling and blood vessels. Slices above the barrel labeling are considered to belong to L2/3, while slices below the barrels are categorized as L5. In each slice, stGtACR2+ neurons were detected using custom MATLAB software, which identifies them based on shape, size, and contrast to the background. Detection in each slice was verified, and if necessary, the detection parameters were manually adjusted to ensure high-quality results. For each cortical column, we counted the stGtACR2+ neurons within a circular area with a radius equal to the distance between two barrels, thereby encompassing the cortical column and its immediate surrounding region. For each cortical compartment (supragranular and infragranular layers), we measured the maximum density of stGtACR2+ neurons across slices within that compartment. This histological analysis provides a single stGtACR2 expression density value for each cortical compartment within each cortical column.

### Analysis of spiking data

Spike sorting was carried out using Kilosort 3, following the standard preprocessing and analysis described in the reference article^40^. All recordings were processed identically using the same thresholds and detection settings. Drift correction was disabled, as it introduced artifacts in recordings with sparse spiking activity. All units assigned by Kilosort3, including unsorted ones, were included in the optogenetic modulation analysis. To avoid contamination by noise, a minimum spike amplitude threshold of 15 arbitrary units was applied to exclude false detections. To identify the center of layer 4 (L4), each electrode channel’s time series was Z-scored, and artifacts (voltage changes exceeding 20 standard deviations) were removed. The depth corresponding to the peak power in the 65–300 Hz range was identified and used as the L4 center. This frequency peak coincided with a current sink in the CSD profile, as illustrated in **Supplementary Figure 3**. The Optogenetic Modulation Index (OMI) was calculated as: OMI=(FR_light-on_+FR_no-light_)/(FR_light-on_−FR_no-light_) where FR indicates the average firing rate across trials.

### Pre-processing and spectral analysis

Pre-processing of the ECoG data was performed as previously described^17^. In brief, for each channel, we applied a common average reference subtraction on the 3kHz signal, subtracting the median of 128 channels. Then a spectral decomposition was performed using a wavelet morse decomposition. Spectral decomposition was performed between 1 Hz and 1.5 kHz with 6 frequency bins per octave. We considered only the real part of the wavelet decomposition. Each frequency band was individually Z-scored based on the statistics of the baseline activity. For baseline statistics we considered all epochs of time with neither light nor whisker stimuli within the previous or following 1 s. We computed standard deviations and average from this baseline activity in a rolling 60s long window, and separately for each wavelet frequency. Z-scoring removes the characteristic power law P ∼ 1/f^a^ fall-off found in brain electrical fields and highlights sensory evoked change in the frequency components. The apparent pre-stimulus ’smearing’ of evoked activity in low-frequency bands is due to the larger time windows and bandwidths characteristic of these frequencies. Finally, for calculation of a full frequency band (e.g. Theta), we averaged the activity over all Z-score whose center frequency falls within the frequency band. We used the following frequency bands: θ (4 - 8 Hz), β (10 - 27 Hz), γ (30 - 57 Hz), Hγ (65 - 170 Hz), uHγ (180 - 450 Hz) and MUA (500 - 1500 Hz). Response to stimuli were measured as +- 5ms around response peak time. Response peak time was computed as maximum Z-score in the high gamma activity band in the 50 ms following the stimulus onset. Data from the extracellular probes was converted to the kilosort format and processed with kilosort3 for spike detection ^40^. Data from both single-units and multi-unit activity were included in the analysis of laminar probe recordings.

### Analysis of whisker tuning and channel selection

One channel was selected to represent each whisker, termed the whisker’s peak-channel. The peak-channel was defined by the user as a manual click at the center of the functional responses observed in the combined Hγ and uHγ bands over the S1 area. Manual selection allowed to avoid unresponsive or broken channels and generated channel selection most coherent with the somatotopic map, as compared to other means of calculation (see **Supplementary Figure 1**). For further analysis, a criterion of responsiveness (average Z-score across all frequencies bands >1) was applied to include only strongly whisker responsive peak-channels. It included 271 out of 324 peak-channels.

### Analysis of optogenetic effect

Optogenetic effect magnitude was quantified either as a difference in the Z-score, or as a ratio. Given that Z-score can approximate null values, it cannot reliably be used to compute ratios. Instead the ratio of suppression was calculated from the wavelet raw power, separately for each frequency band.

Grouping of the peak-channels for population analysis (**Fig. 3**) was made using the same reasoning as for the statistical model. Inclusion in the group requires putative suppression in the defined layers. The suppression of firing rate in a layer depends on the necessary interaction between the opsin expression and the presence of optogenetic light. We used the following criteria of optogenetic light exposure and laminar stGtACR2 expression to constitute the four groups: **Control group**: no expression of stGtACR2 in the whole animal. **L2/3 suppression group**: criterion (1) [stGtACR2 expression in L2/3 x λ_spatia_] is above the 70 percentile across all peak-channels, and criterion (2) not in the L23 & L5 suppression group below. **L5 suppression group**: criterion [stGtACR2 expression in L5 x light level] is above the 70 percentile across all peak-channels. **L2/3 & L5 suppression group**: criterion stGtACR2 expression in L5 x stGtACR2 expression in L2/3 x light level is above the 90 percentile across all peak-channels, non overlapping with the other groups.

We used a mixed effect model to assess the contribution of each layer to optogenetic modulation of a peak-channel’s average CSEP (Yu et al 2022). The dependent variable, ΔZsc, is the difference in sensory-evoked activity between light and no-light conditions for a given peak-channel at a given optogenetic light intensity (n = 717 suppression data points from 153 peak-channels in 17 recordings, multiple intensities per peak-channel). For a given peak-channel, ΔZsc is fitted from the putative suppression of AP activity in the different layers as:

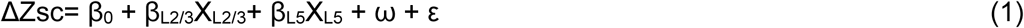

Here:

- ω represents a nested random effect (whisker identity in the recording).
- ε is the residual error term.
- β_L2/3_ and β_L5_ are the model parameters (i.e. weights) estimating the contribution of each layer to ΔZsc.
- X_L23_ (in L2/3) and X_L5_ (in L5) represent the putative suppression of firing rate in pyramidal neurons in each layer. The putative suppression of pyramidal neurons depends on the interaction between the exposition to optogenetic light at the column underneath (λ_L2/3_ and λ_L5_) and the laminar opsin expression in L2/3 (stGtACR2_L23_) or in L5 (stGtACR2_L5_).

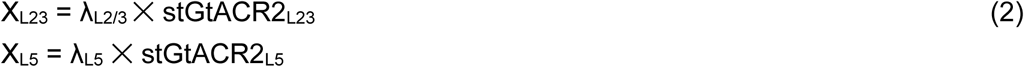

Where:

- stGtACR2_L23_ and stGtACR2_L5_ are estimated as the densities of stGtACR2-expressing neurons per mm2 in L2/3 and L5 respectively (see histology section for details)
- λ_L2/3_ and λ_L5_ are unit-free numbers estimating the light reaching neurons in L2/3 and L5, computed as:

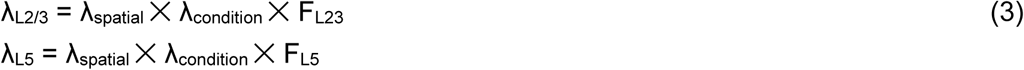

λ_spatial_ is measured as the artifact produced by light in the peak-channel. As light was delivered as a square pulse, it created a sizable artifact at light onset. λ_spatial_ was computed as the Pearson’s R between the optogenetic pulse first derivative and the raw voltage first derivative yielding an area of light clearly localized and matching the position of the optical fiber. λ_condition_ is the light magnitude delivered on the condition (0, 0.9, 1.9, 3.5, 6.4, 11.0, or 18 mW). F_L23_ and F_L5_ are the irradiance fraction of the light reaching the mean depth of the layer (F_L23_, = 0.39 at 200 µm for L23 and F_L5_ = 0.19 at 550 µm for L5) as predicted by the stanford scale (https://web.stanford.edu/group/dlab/cgi-bin/graph/chart.php). Finally, as we observed saturation in the reduction of neuronal firing rate as a function of light intensity, we applied a ceiling to both λ_L2/3_ and λ_L5_ at a value of 0.0257 (arb. units), corresponding to 3.5 mW of light intensity in the spatially most illuminated channel. This is the level above which we observe no increase in suppression with increasing light intensity. To compute the proportion of variance explained, we used a leave-one-out procedure and calculated the coefficient of determination R² = 1-RSS/TSS with RSS is the residual sum of squared error and TSS the total sum of squared error. To estimate statistical significance of the difference between β_L2/3_ and β_L5_, we created a null distribution by shuffling randomly the peak-channels order in the matrices X_L2/3_ and X_L5_ and refitting the model 10^5^ times. The experimentally observed difference was compared to the null distribution, and a statistical p-value computed as p =2 x (1-|percentile|), to correspond to a two-sided test.

## Supplementary information

**Supplementary Table 1.** Group by layer expression.

| Group | Number of peak-channels | Animals breed | Criteria (Histologically confirmed) |
| --- | --- | --- | --- |
| ACR2 - Control | n = 63 | N = 3 WT, 3Drd3, 2 Rbp4 | no expression |
| ACR2+ L2/3 | n = 25 | N = 6 Drd3 | >70<sup>th</sup> percentiles L2/3, and not in ACR2+ L2/3 x L5 group |
| ACR2+ L5 | n = 35 | N = 6 Rbp4 | >70<sup>th</sup> percentiles L5 |
| ACR2+ L2/3 x L5 | N = 15 | N = 4 Drd3 | >90<sup>th</sup> percentile L5 x L2/3 |

**Supplementary Table 2.** Group by layer expression.

| *Metric* | *Theta* | *Beta* | *gamma* | *Hγ* | *uHγ* | *MUA* |
| --- | --- | --- | --- | --- | --- | --- |
| *Ratio BL5/BL23* | 0.50 | 0.49 | 0.90 | 2.06 | 1.66 | 1.00 |
| *Percentile BL23 -BL5* | 99.95 | 99.86 | 68.57 | 00.01 | 00.64 | 48.65 |
| *Two-sided p-value* | 0.0010 | 0.0028 | 0.6286 | 0.0002 | 0.0128 | 0.9730 |
| *Adjusted p-values (Benjamini-Hochberg)* | **<br>0.0023 | **<br>0.0049 | 0.7334 | ***<br>0.0007 | *<br>0.0179 | 0.9730 |
| *Coefficient of determination R<sup>2</sup> (held out)* | 0.43 | 0.35 | 0.26 | 0.34 | 0.28 | 0.27 |

**Supplementary Figure 1:**
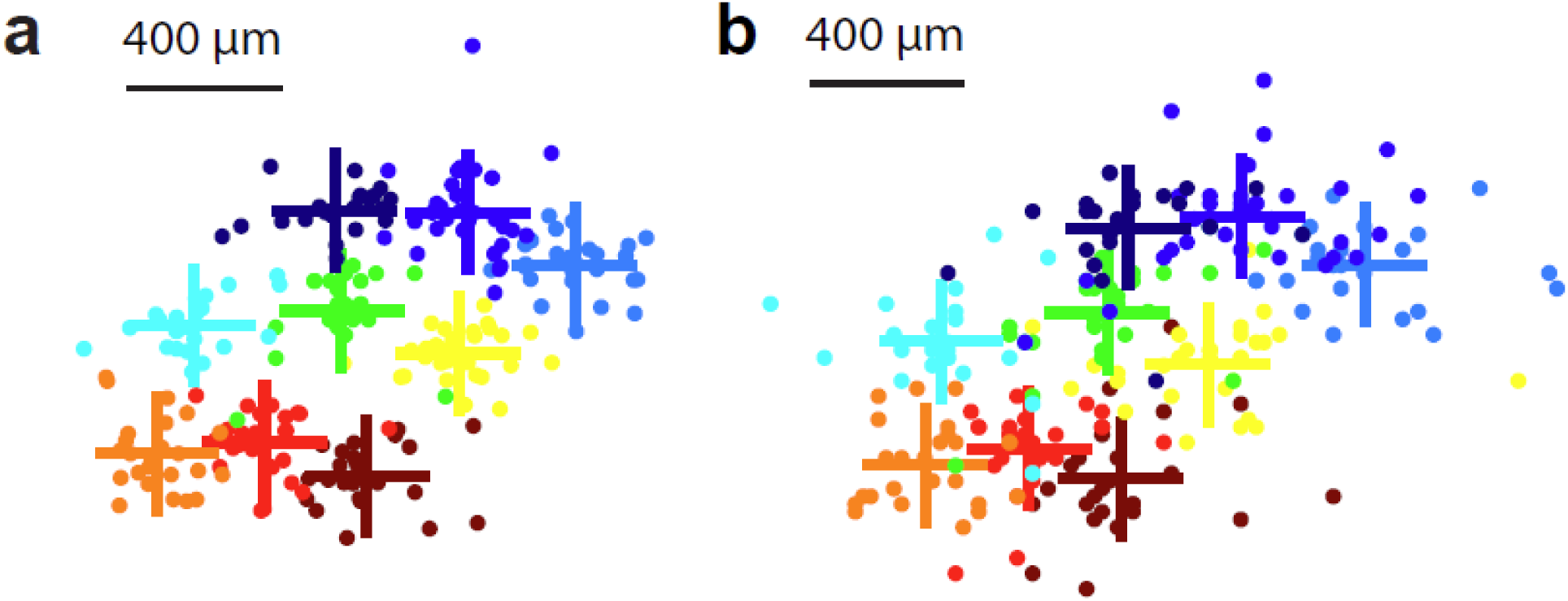
Peak Channel selection. Manual and automated methods for peak-channel selection both show clear somatotopy. **a**, Manual selection which was used for all analyses using peak-channels throughout the study. **b**, Spatial peak of Hγ activity also displays robust somatotopy, with individual points more scattered than in the manual method. This difference may arise from electrode properties or varying contact with the underlying tissue. Both maps were aligned across animals by recursively minimizing the Euclidean distance between all whiskers and their averaged position in the pattern, with a resolution of 20 µm (interpolated at 10x).

**Supplementary Figure 2:**
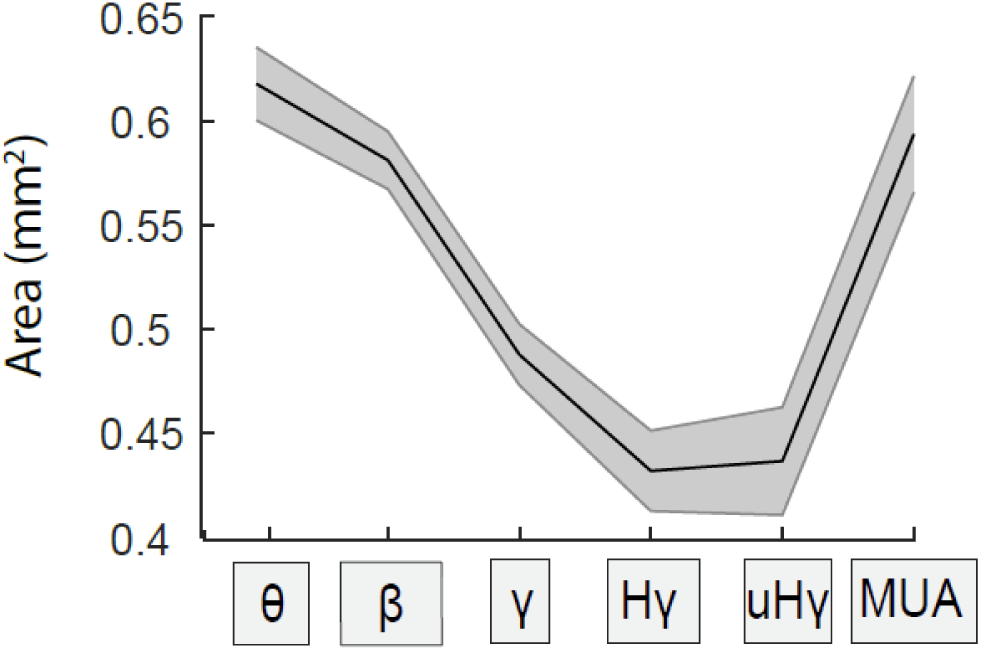
Spatial spread of stimulus-evoked activity across different frequency bands in CSEPs. Spread was quantified as the area where responses exceeded half the peak activity in the endogenous frequency, averaged across responsive peak-channels. Error shades represent sem. This pattern is qualitatively similar to observations in the rat auditory cortex. The rise in MUA may, in part, reflect a reduced signal magnitude in that frequency range.

**Supplementary Figure 3.**
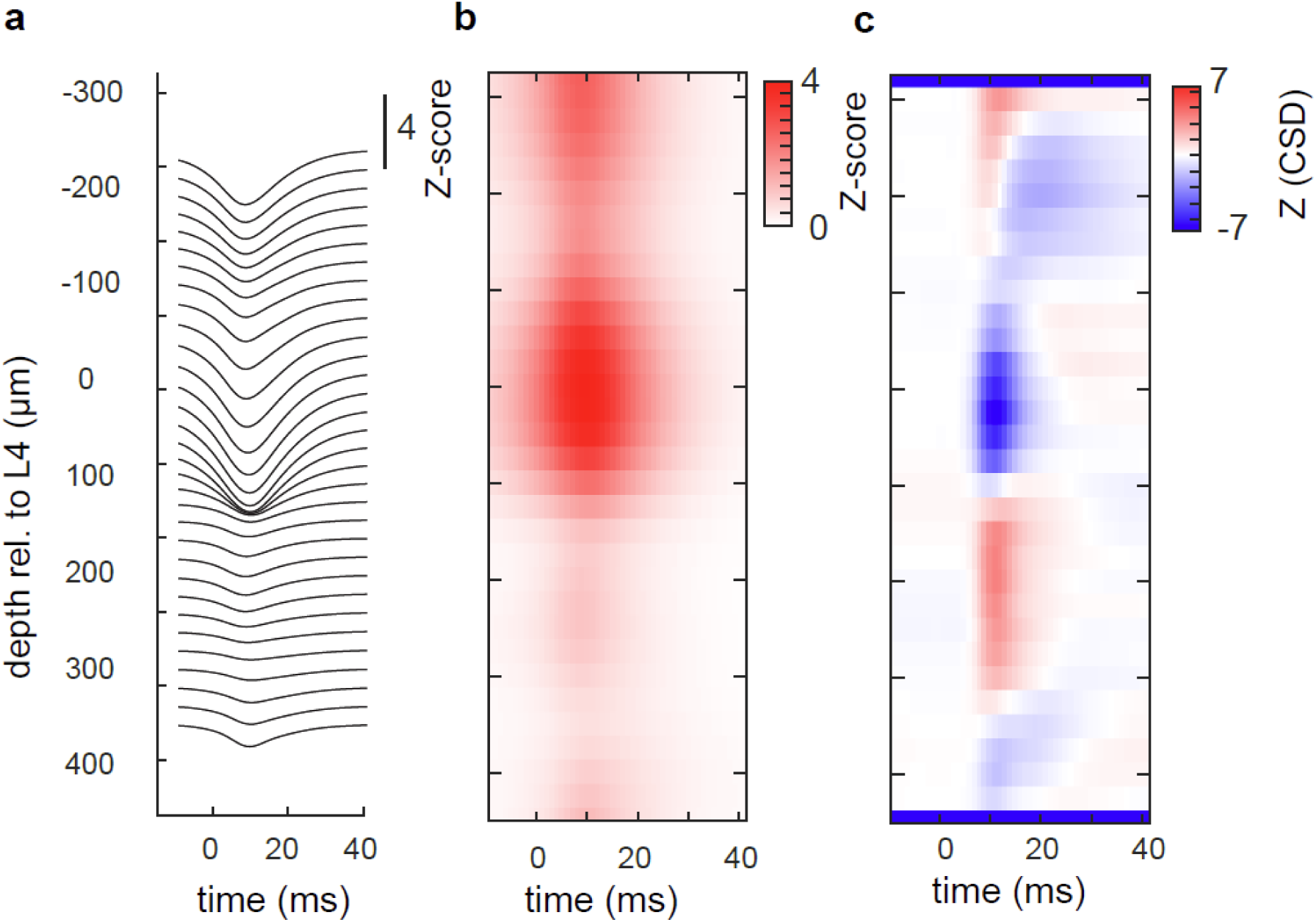
Localization of layer 4 center from laminar recordings. **a**, Peri-stimulus voltage traces from 32 channels in an example electrode penetration, averaged across trials for the preferred whisker. Signals were Z-scored and bandpass filtered between 1–300 Hz. **b**, Corresponding high-frequency activity used to localize the center of layer 4. The depth with maximal energy in the 10–20 ms window post-stimulation was identified as the L4 center. **c**, Corresponding current source density (CSD) analysis, computed as the second spatial derivative of the voltage signal. Traces were filtered (1– 300 Hz) and smoothed in both spatial and temporal dimensions (σ = 2) prior to CSD computation.

